# Structural insights on the ligand selectivity of the mouse trace amine-associated receptor TAAR7e

**DOI:** 10.1101/2025.08.25.669582

**Authors:** Yingjian Liu, Junhong He, Jiahui Sun, Chen Zhang, Hanyi Zhuang, Peipei Shi, Weihong Liu

**Affiliations:** Intelligent Perception Lab, Hanwang Technology Co., Ltd, Beijing 100193, China

**Author notes:** Correspondence author. E-mail addresses (Y. Liu), (W. Liu). These authors contributed equally to this work (Yingjian Liu, Junhong He, Jiahui Sun).

**Keywords:** GPCR, G protein-coupled receptor, SAR, structure-activity relationship, TAAR, trace amine-associated receptor

## Abstract

Trace amine-associated receptors (TAARs), a subclass of G protein-coupled receptors involved in detecting volatile amines, are co-expressed with odorant receptors in the main olfactory epithelium. Although many behaviorally relevant amines have been identified for olfactory TAARs, the molecular mechanisms underlying their selectivity for amines with varying carbon chain lengths and degrees of amine substitution remain unclear. In this study, we used mTAAR7e as a model to investigate these mechanisms. Homology modeling revealed a conserved ligand-binding pocket, supported by sequence-structure covariance analysis across TAARs. Structure-activity relationship profiling revealed key chemical determinants integral to mTAAR7e-mediated odorant recognition and provided, for the first time, a structural explanation for its selective preference toward longer carbon chains and tertiary amines. Computationally predicted interactions between mTAAR7e and the representative ligand *N,N*-dimethylcyclohexylamine (DMCHA) were validated through site-directed mutagenesis. Furthermore, conformational dynamics of mTAAR7e during receptor activation were characterized, providing insights into activation-related structural rearrangements. Together, these findings offer novel insights into the molecular logic of TAAR ligand selectivity and may advance our understanding of how TAARs mediate both olfactory and systemic aminergic signaling.

## 1. Introduction

Trace amine-associated receptors (TAARs) are a group of G protein-coupled receptors that are responsive to trace and volatile amines [1]. With a total of 15 functional TAAR genes in the mouse genome and 6 in the human genome, the olfactory TAARs (TAAR subfamilies 2-9) are expressed alongside olfactory receptors (ORs) in the mammalian olfactory epithelium [1-3] and ectopically [4]. While olfactory TAARs may serve as a basis for ecological questions such as predator-prey interactions, mate selection, nutrient, and habitat choices, the physiological roles of TAARs expressed in non-olfactory tissues promise immense therapeutic potentials. In parallel to the deorphanization effort of ORs, past studies have revealed ligands for a number of TAARs. As the name suggests, the ligands of TAARs are aminergic; yet diverse structures are seen within the TAAR ligand repertoire [3, 5, 6]. Ranging from primary to tertiary, aliphatic to aromatic, monoamines to polyamines, the biological functions of most of these volatile ligands are olfactory cues relevant to various aspects of animal behaviors. This is with the exception of TAAR1, which is mainly expressed in the brain and is responsible for the activity against endogenous brain-derived trace amines and certain psychotropic agents [2].

Previous studies have examined the structural features constituting the ligand selectivity of TAARs. The length of carbon chain directly affects ligand size and is one of the determining factors in eliciting a response. The effect of carbon chain length was assessed in the zebrafish TAAR13c, which only responds to 5-carbon, 6-carbon, and 7-carbon medium-chain diamines, with a preference for the 5-carbon and 7-carbon odd-numbered chains, possibly as a result of the orientation of the two amine groups [7].

Positively charged amine groups require either an aspartic acid or a glutamic acid for ligand coordination. Li *et al*. suggested a possible structural basis for the evolution of the selectivity for the number of amines in TAAR ligands through the study of the diamine cadaverine receptor TAAR13c. The involvement of two remote transmembrane aspartic acids, Asp^3.32^ and Asp^5.42^, forming a di-cation recognition motif in the receptor, is required for diamine response whereas the presence of either Asp^3.32^ or Asp^5.42^ confers response to primary or tertiary monoamines [8]. A similar diamine binding pocket with Asp^3.32^ and Asp^5.43^ is seen in the mouse and human TAAR6s and TAAR8s but whether these receptors are responsive to diamines is not known [8, 9]. One exception to the rule above is TAAR9, which only has Asp^3.32^ and can recognize ligands with one, two, and three amine groups [6], pointing to the possibility of the existence of yet another cation recognition site.

Finally, it has been proposed that nine mammalian TAAR subfamilies can be further classified into two clades based on the types of amines they detect, with TAAR1, TAAR2, TAAR3, and TAAR4 being specific for primary amines and TAAR5, TAAR6, TAAR7, TAAR8 and TAAR9 for tertiary amines [5]. The dichotomy by ligand type is also corroborated by sequence similarities and by phylogenetic analysis, in which proteins in TAAR1-4 cluster distinctively from those in TAAR5-9, the latter of which evolved more recently and under positive selection, allowing for a higher rate of gene duplication events [10]. However, the molecular basis for the preference of more substituted amines in TAAR5-9 remains unknown.

While TAAR7 is a pseudogene in the human genome, the mouse TAAR7s underwent lineage-specific expansion to contain five members sharing high sequence identities yet exhibiting different expression locales and diverse response profiles against various tertiary amines. Specifically, mTAAR7e recognizes a variety of amine odorants, including primary, secondary, and tertiary amines, to generate distinct physiological responses. Various screening efforts revealed that its agonists include 5-methoxy-*N,N*-dimethyltryptamine, a urinary tryptophan metabolite, *N,N*-dimethyl-2-phenylethylamine, *N,N*-dimethylbutylamine (DMBA), *N,N*-dimethyloctylamine (DMOA), 2-methyl-1-pyrroline, heptylamine, octylamine (OA), etc. [5, 6]. Both TAAR7e and TAAR7f are expressed exclusively in the dorsal portion of the olfactory epithelium [11] and the amino acid sequence identity between the two receptors is 93%. Ferrero *et al*. showed that each of the two receptors preferentially binds to certain ligands as a result of two adjacent amino acid side chains in the ligand-binding pocket in TM3. Changing the amino acids at 132^3.37^ and 133^3.38^ of TAAR7e to those of TAAR7f resulted in a response profile similar to that of TAAR7f and vice versa. Homology modeling (HM) revealed that the bulky side chain of tyrosine 132 in TAAR7f hinders interactions with large ligands [5]. This shows that the broadened receptive range of TAARs over the course of evolution is a result of gene duplication followed by mutations in ligand-binding pockets.

The mode of interaction and the tuning of the TAARs are of interest to both the scientific and medical communities. Recently, several cryo-EM structures of TAARs were resolved, including mTAAR9 [12], mTAAR7f [13], and TAAR1 [14-16]. The structures of both chemosensory TAARs, mTAAR9 and mTAAR7f, were determined against the tertiary amine ligand, dimethylcyclohexylamine (DMCHA). Although the pose of the ligand was different between the two structures, it was observed that the receptors bind to the tertiary amine through a charge-charge interaction with the carboxylate group of the aspartic acid residue Asp117^3.32^ in mTAAR9 [12] and Asp127^3.32^ in mTAAR7f [13]. The same aspartic acid was also implicated in the binding of mTAAR9 to spermidine and β-phenylethylamine and in the binding of mTAAR1 and hTAAR1 to its ligands [14-16].

Here we elucidate the structure-activity relationship (SAR) of mTAAR7e by using a series of structurally related ligands. By constructing a homology model using the closely related mTAAR7f structure, we home in on the TAAR7e binding pocket and identify the structural basis for tertiary amine preference, which, along with conformational dynamics studies, constitutes a starting point to design a computational strategy for elucidating the ligand selectivity of all TAARs.

## 2. Experimental section

### 2.1. Luciferase assay

Luciferase assay was performed using Dual-Glo™ Luciferase Assay System (Promega Biotech Co., Ltd., Beijing, China) as previously described [17]. HEK293T-derived cells were plated on 96-well plates (Corning Inc., Kennebunk, ME, USA). For each 96-well plate, we transfected 1 μg of CRE-Luciferase, 0.5 μg of pRL-SV40, 0.5 μg of TAAR, 0.5 μg of RTP1S, and 0.25 μg of M3R using the transfection reagent Lipofectamine2000 (Invitrogen, Life Technologies Corp., Carlsbad, CA, USA). Twenty-four hours post transfection, we removed the transfection media and rinsed the cells with 50 μL PBS buffer (pH 7.4) per well. Then we immediately replaced the PBS buffer with 25 μL TAAR ligands diluted in CD293 medium (Gibco Brand, ThermoFisher Biochemical Products Co., Ltd., Beijing, China). Four hours post stimulation, Firefly luciferase (FL) and *Renilla* luciferase (RL) acttiffies were measured following manufacturer’s protocols, using a Synergy H1 plate reader (BioTek Instruments, Winooski, VT, USA). We calculated normalized luciferase activity using the formula: (FL/RL_X_ ⍰ FL/RL_XC_) / (FL/RL_max_ ⍰ FL/RL_maxC_), where FL/RL_X_ is the response of the wild-type or a mutant TAAR to a test sample; FL/RL_XC_ is the basal value of the same receptor to a blank control; FL/RL_max_ is the maximum response of the wild-type or a mutant TAAR to the same or another ligand, and FL/RL_maxC_ is the basal value of the receptor with the maximum response to a blank control.

### 2.2. Homology modeling and docking

HM structure of mTAAR7e was conducted using Swiss Model [18] based on a high-resolution cryo-EM structure of mTAAR7f (PDB ID: 8PM2) and the quality of models were measured by Ramachandran plot [19].

All docking calculations were carried out by AutoDock Vina [20]. ADFR protocol in AutoDock Vina was implemented for the preprocess of receptors. During preprocessing, protein structural problems such as missing excitation structures and steric clashes were identified and corrected. Then the receptor was optimized by applying the OPLS_2005 force field [21]. The grid box utilized in docking was determined by the atomic center of the ligand in the mTAAR7f, and the box size was set up to 15 Å. The ligand conformations were optimized and generated using the Python package Meeko before docking and scoring by autogrid4 [22].

### 2.3. Molecular dynamic simulation

All MD simulations were conducted using the GROMACS package (version 2020)[23] with the ff14SB force field [24], starting from docking structures. The complex system was prepared by pdb4amber tool in AMBERTOOLS 2023 [25] and then solvated using the TIP3P water model. As the topology of the complex system was established, the heavy atoms of the system were minimized through a three-step minimization process, with individual position restraints set at 5, 3, and 0□ kcal/(mol·Å^2^), followed by a 1-ns heating step, which raising the temperature from 0 K to 300 K in NVT ensemble with a Nosé-Hoover thermostat [26]. Subsequently, a 10-ns equilibration process was conducted, bringing the system to a relatively stable state. During the heating and equilibration steps, the position restraints was changed to 3□kcal/(mol·Å^2^). The last frame of equilibration was used as the initial conformation for three production runs with random seeds.

### 2.4. Molecular Dynamic analysis

#### 2.4.1. Ligand-receptor interactions

Contact frequencies were calculated using the Python package named “get contacts” (https://getcontacts.github.io/), and the following contacts between specific atoms of ligands and residues of receptors were counted: hydrogen bonds, hydrophobic and van der Waals interactions.

#### 2.4.2. RMSD

The backbone atoms of receptors were used to calculate the Root Mean Square Deviation (RMSD) by gmx rms (GROMACS package 2019) [23] function to determine whether simulations were stable.

#### 2.4.3. RMSF

The Root Mean Square Fluctuation (RMSF) analysis was performed on the C_α_ atoms of the receptor using the MDanalysis [27] Python package. The transmembrane backbones and extracellular loops were labeled on RMSF plot as follows: 48-68 (TM1), 84-104 (TM2), 122-143 (TM3), 167-187 (TM4), 213-233 (TM5), 275-295 (TM6), 310-333 (TM7), 188-212 (ECL2), 296-309 (ECL3).

#### 2.4.4. Free energy calculation

The Free energy was calculated by Molecular Mechanics Poisson-Boltzmann Surface Area (MMPBSA) [28] with final 100ns snapshots from an ensemble of conformations.

#### 2.4.5. Distance of residue

To calculate the distance between two residues, specific atoms within each residue were selected and then the gmx distance tool was utilized to calculate these measurements for all frames in the simulations. The reference distance in the initial conformation of the complex system was determined using the measurement tool in PyMOL [29].

## 3. Results and discusssion

### 3.1. Overall architecture of mTAAR7e and the orthosteric binding site

A homology model of mTAAR7e was generated using the published mTAAR7f structure as a template [13]. Given the high sequence identity and template quality, most residues adopted sterically favorable conformations, as validated by Ramachandran plot analysis (SI Fig. S1). Manual inspection of the remaining residues revealed they were primarily localized in loop regions, which are unlikely to affect the orthosteric binding site (OBS) within the helix bundle.

To better understand the binding mechanism of mTAAR7e, we conducted a sequence conservation analysis of the OBS residues (within 3.9 Å of each ligand) in the mTAAR7e-ligand complex structure obtained from molecular docking, the TAARs and OR51E2 structures resolved by cryo-EM experiments. This analysis revealed the presence of a conserved DWY motif (Asp127^3.32^, Trp286^6.48^, Tyr316^7.42^) and a functionally distinct residue substitution at position 5.43 in TAARs (Fig. 1a). This phenomenon was also confirmed by extensive sequence alignment of mTAARs (Fig. 1d-e). The olfactory receptor OR51E2 exhibits limited structural similarity and minimal sequence conservation compared to TAARs [30], reflecting fundamental mechanistic differences in odorant recognition between human ORs and mouse TAARs. Notably, ligand-binding residues in OR51E2 overlap with TAARs only at positions 5.43 and 3.33, which are hypothesized to play roles in receptor activation pathways.

**Fig. 1.**
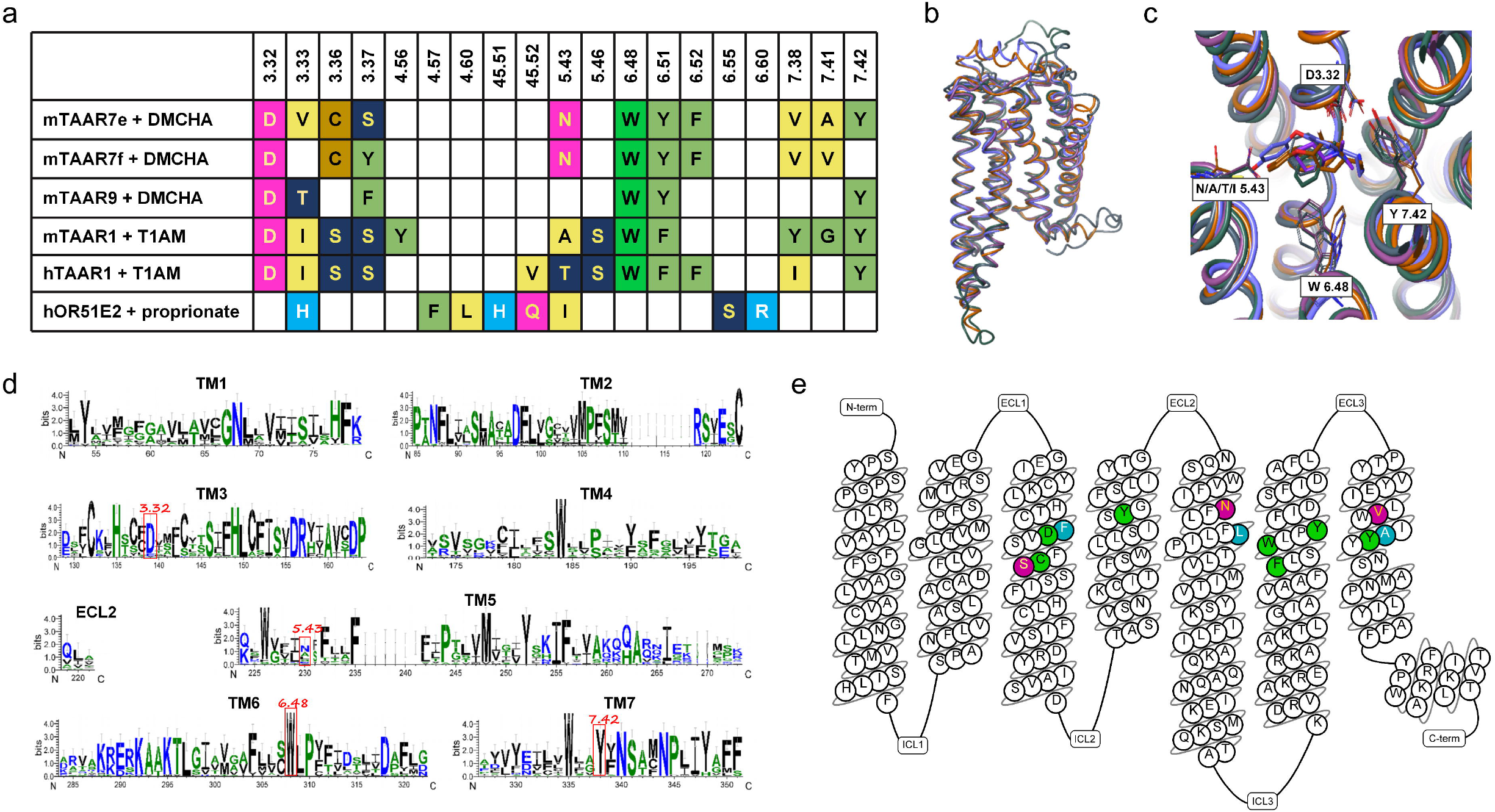
The orthosteric binding pocket of mTAAR7e. (a) Residues in the OBS within 3.9 Å of the ligand of mTAAR7e, mTAAR7f, mTAAR9, mTAAR1, hTAAR1, and hOR51E2 (PDB IDs: mTAAR7f, 8PM2; mTAAR9, 8ITF; mTAAR1, 8JLJ; hTAAR1, 8JLN; OR51E2, 8F76). (b) Superposition of the agonist-bound structures of mTAAR7e (pine), mTAAR7f (violet), mTAAR9 (gray), mTAAR1 (pale blue), and hTAAR1 (orange). (c) Comparison of side chain positions in the receptor alignments from the extramembrane perspective. (d) Multiple sequence alignment of mTAARs. (e) The snake plot of mTAAR7e highlighting the OBS, with green residues representing those conserved among mTAARs, blue residues indicating differences from mTAAR7f but conserved across other mTAARs, and purple residues corresponding to divergent residues within the mTAAR family.

Building on these sequence insights, comparative analysis of the mTAAR7e model with experimentally resolved TAAR structures revealed high structural conservation in the transmembrane (TM) region, with an average RMSD of less than 1 Å (Fig. 1b), providing a robust foundation for subsequent OBS conformation analysis (Fig. 1c). In DWY motif, Asp127^3.32^ occupies the same position on TM3 and the carboxylic acid side chain is located near the nitrogen atom of the ligand, while the positions of Trp286^6.48^ and Tyr316^7.42^ in mTAAR7e, mTAAR7f, and mTAAR9 are also highly similar. In contrast, mTAAR1 and hTAAR1 exhibited rotation of Trp286^6.48^ towards TM7 due to steric hindrance from isoleucine at position 3.40 [13].

### 3.2. Ligand selectivity of mTAAR7e

A previous report has shown that DMBA may be a ligand of mTAAR7e [6]. We confirmed this result with a full dose-response curve (Fig. 2a). We estimated the logEC_50_ to be around −5.61 (∼2.4 μM) (Table 1). To examine the ligand selectivity of mTAAR7e, several structurally related ligands were tested in an attempt to further depict the ligand-binding pocket and to discern the functional groups responsible for receptor-ligand interaction. In comparison to DMBA, DMOA showed increased efficacy and a significantly increased potency (Fig. 2b), boasting a logEC_50_ of around −6.57 (∼0.27 μM) (Table 1). We note that the EC_50_ values obtained for DMBA and DMOA are more than 30-fold and 270-fold lower than the previously reported values, respectively, reflecting the sensitivity of the cell system used this study. In contrast to the substituted amines, butylamine, with hydrogens replacing the methyl groups, showed a significant reduction in potency, rendering the receptor almost inactive (SI Fig. S2a). This indicates the methyl groups may be important in anchoring to the receptor. Similarly, OA exhibited reduced receptor activity (Fig. 2c), recapitulating the importance of the dimethyl group at the ligand-receptor interface. We also decreased the length of the carbon chain by using trimethylamine, a known ligand of some of the other members of the TAAR family. We found only a minimal level of activation (SI Fig. S2b), indicating that decreasing the length of the carbon chain adversely affect receptor efficacy. To investigate the effect of steric hinderance in the binding pocket, we included DMCHA. We found this compound showed decreased potency compared to DMBA (Fig. 2d). In summary, profiling activities of mTAAR7e toward various structurally related ligands shows that the ligand-binding pocket of mTAAR7e is amenable to longer-chain aliphatic carbon backbone and with a strong preference for the terminal dimethyl group.

**Table 1.**
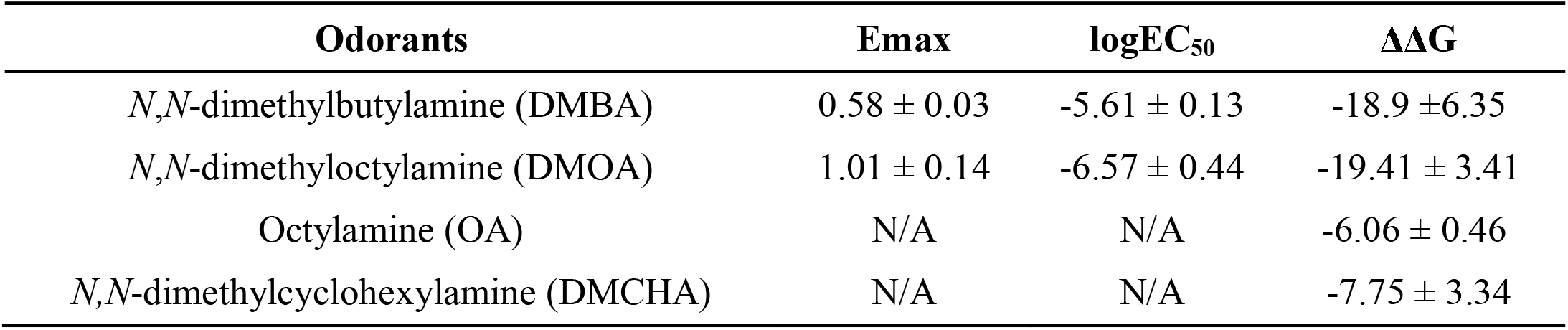
The E_max_, logEC_50_, and ΔΔG of wild-type mTAAR7e on selected odorants.

**Fig. 2.**
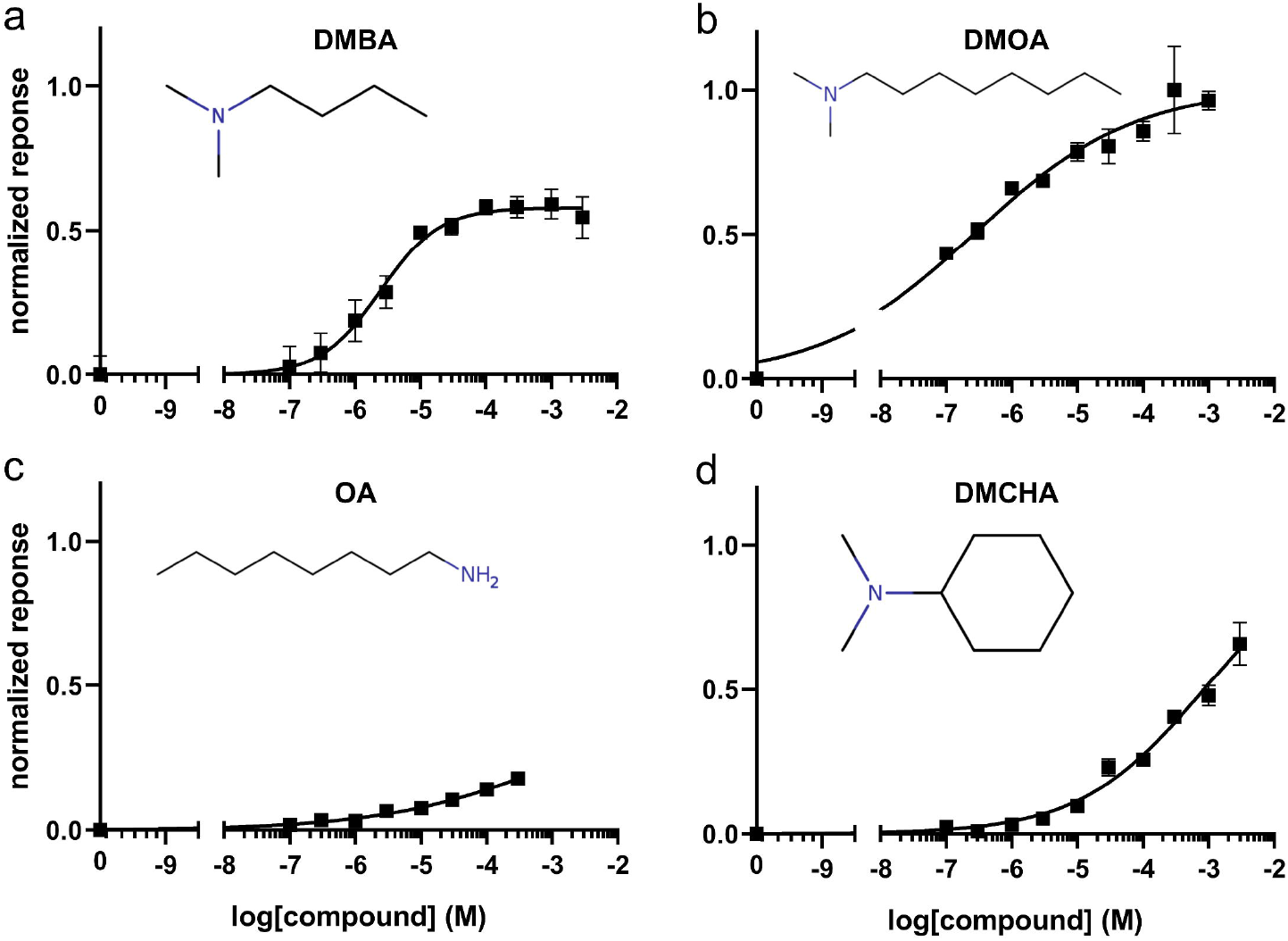
Responses of mTAAR7e to selected compounds. All responses are normalized to the highest response of DMOA. *N* = 4.

Experimental results have established the binding affinity hierarchy of mTAAR7e as DMOA > DMBA > OA, with computational analysis of these complexes illuminating the underlying selectivity mechanism (SI Fig. S3a-c). MMPBSA free energy calculations from MD simulations highlighted residues with significant contributions (|ΔΔG| > 1 kcal/mol) (Fig. 3a-c), where Asp127^3.32^ consistently emerged as the primary contributor across all ligands, underscoring its critical role in binding. DMOA and DMBA exhibited similar energy contribution patterns whereas the absence of the dimethyl group in OA led to distinct interactions. Structural visualization of the binding pockets for DMOA and DMBA (Fig. 3d) revealed that DMOA#x2019;s longer carbon chain conferred superior steric complementarity, explaining its higher affinity. Decomposition of free energy differences between DMOA and OA complexes (Fig. 3e) identified Trp286^6.48^ and Tyr316^7.42^ as key divergent residues, driven primarily by electrostatic (ΔE_EEL_) and van der Waals (ΔE_VDW_) interactions, respectively. Previous evolutionary analyses provide background for this selectivity for amine compounds: mTAAR subfamilies diverged under distinct selective pressures, with mTAAR5-9 (responsive to tertiary amines like DMOA/DMBA) undergoing gene duplication and positive selection, while mTAAR1-4 (responsive to primary amines like OA) remained more evolutionarily conserved [10]. Hydrophobic comparisons of mTAAR1 and mTAAR7e revealed significant residue variations within the OBS, notably at positions 5.43, 6.51, 6.52, and 7.38—among which position 5.43 showed the most dramatic shift (Fig. 3f), transitioning from a hydrophobic residue (Ala/Leu) in primary amine-sensing mTAARs to hydrophilic residue (Asp/Asn/Ser) in mTAAR7e and most tertiary amine-sensing mTAARs (except mTAAR9) (Fig. 3g). Given the critical role of Asp127^3.32^ in ligand recognition, we speculate that the structural differences in mTAARs’ amine recognition capabilities are primarily due to residue 5.43 at the top of the pocket. In mTAAR5-9, this position is occupied by hydrophilic residue (Asp/Asn/Ser) and Cys, which are more likely to form hydrogen bonds with primary amines, thereby preventing primary amines from entering the binding pocket and interacting with Asp127^3.32^. The hydrophobicity and charge distribution of the DMOA binding pocket were also demonstrated to support this hypothesis (Fig. 3h-i).

**Fig. 3.**
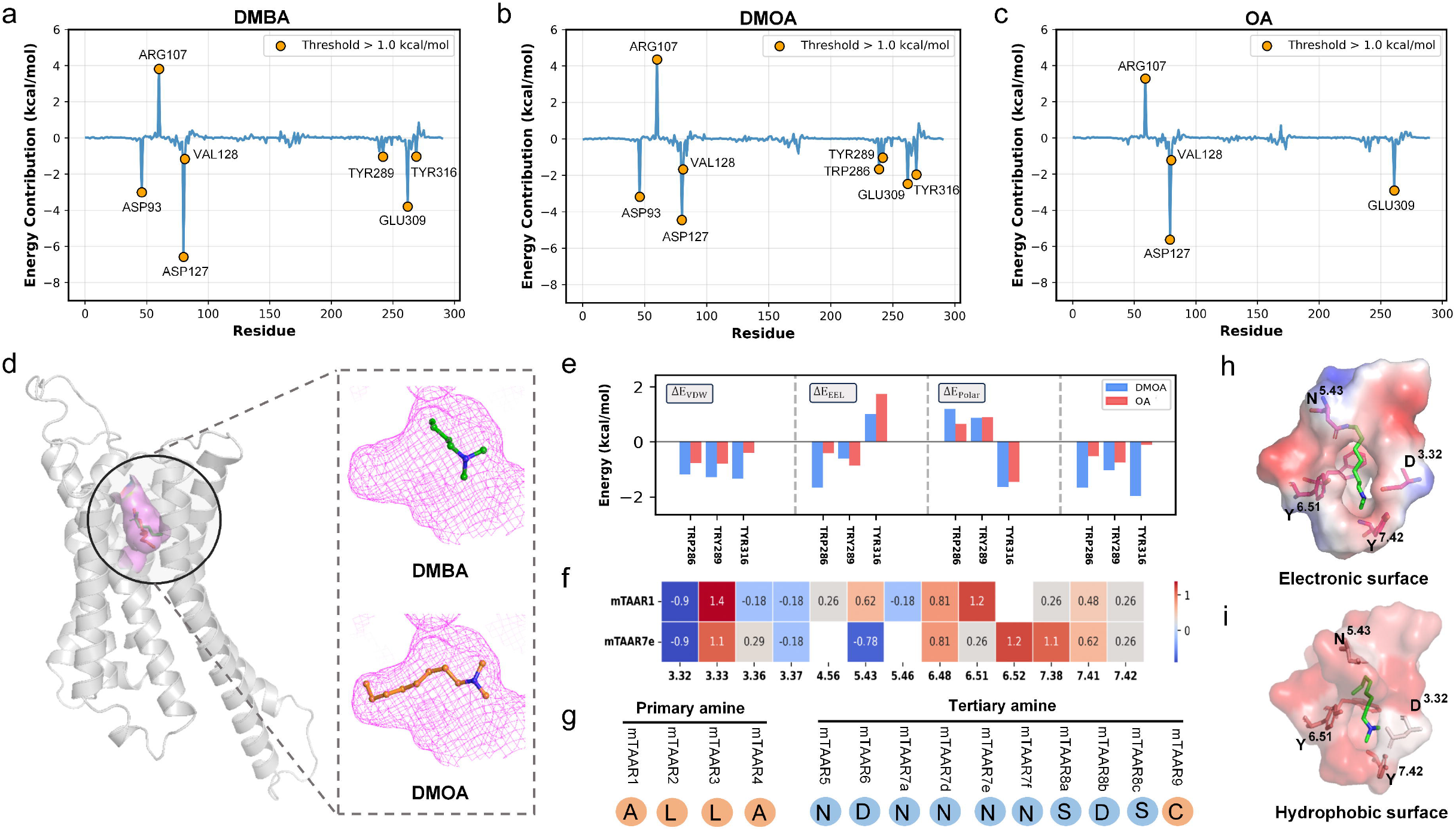
Computational analysis on ligand selectivity. (a-c) MMPBSA energy contribution analysis of residues in mTAAR7e binding to DMBA, DMOA, and OA from MD simulations, with residues showing absolute free energy values > 1 kcal/mol highlighted. (d) Binding pockets of mTAAR7e with DMBA and DMOA. (e) Key differences in binding pocket free energy contributions from residues interacting with DMOA versus OA. (f) Hydrophobic comparisons of residues in the OBS within 3.9 Å of the ligand of mTAAR7e and mTAAR1. (g) Hydrophobic comparisons of residue 5.43 in all mTAARs. (h-i) Hydrophobic and electronic surfaces of the binding pocket between DMOA and mTAAR7e.

### 3.3. Binding interactions between mTAAR7e and DMCHA

Leveraging the structural insights garnered from experiments, we delved deeper into the molecular determinants of ligand interaction with mTAAR7e by calculations, with DMCHA serving as an illustrative example of the binding mechanism (Fig. 4a). We found that the binding pocket consists of 11 residues: 4 hydrophobic (Val128^3.33^, CYS131^3.36^, Val312^7.38^, Ala315^7.41^), 4 aromatic (Trp286^6.48^, Tyr289^6.51^, Phe290^6.52^, Tyr316^7.42^), and 3 polar (Asp127^3.32^, Ser132^3.37^, Asn217^5.43^). Key stabilizing interactions include: a hydrogen bond with Asp127^3.32^, π-cation interaction with Tyr289^6.51^, and π-π stacking with Trp286^6.48^ (Fig. 4d). Three rounds of 300-ns MD simulations on the complex system (SI Fig. S4, SI Table S1) showed a strong polar interaction between the charges on Asp127^3.32^ and DMCHA’s tertiary amine, with the relative distance stabilizing at 2–4 Å (Fig. 4b), while the distance between the benzene ring centers of DMCHA and Trp286^6.48^ stabilized at 4–6 Å, highlighting the importance of these interactions (Fig. 4c). Additionally, the contact frequency heatmap was analyzed, revealing high interaction frequencies between the odorant atoms and residues Asp127^3.32^, CYS131^3.36^, Trp286^6.48^, and Tyr289^6.51^ within the OBS (Fig. 4e). Beyond the established hydrogen bonding, van der Waals contacts were consistently observed, demonstrating complementary non-covalent stabilization mechanisms. The low RMSF values of key residues in the binding pocket were also demonstrated, confirming our conclusions (Fig. 4f).

**Fig. 4.**
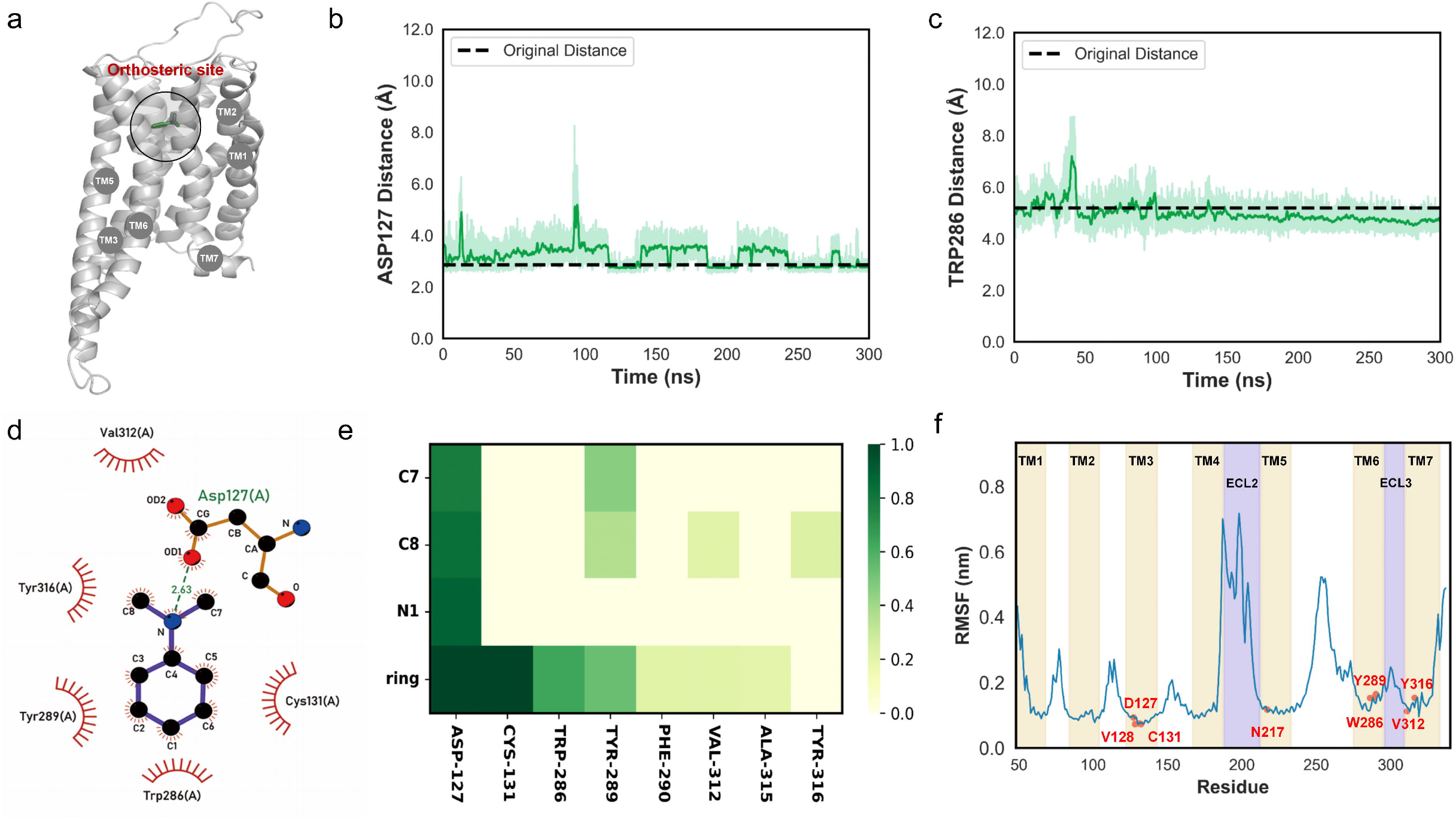
Binding interactions and dynamic characteristics of DMCHA with mTAAR7e. (a) Overall structure of mTAAR7e, the position of OBS is circled. (b) Distance fluctuations between the nitrogen atom of DMCHA and the Asp127 residue during MD simulations. (c) Distance fluctuations between the aromatic ring of DMCHA and the Trp286 residue during MD simulations. (d) Binding pose of DMCHA with mTAAR7e (green for hydrogen bond, red for π-cation, and blue for π-π stacking). (e) Heatmap depicting contact frequencies of interactions between mTAAR7e binding site residues and DMCHA atoms (contact frequency threshold set at 20%). (f) RMSF of MD simulations, the residues in OBS are labeled as red points.

The binding patterns identified through computational modeling were further supported by mutagenesis experiments. In most instances, introducing mutations at key residues within the OBS significantly reduced mTAAR functional activity, as demonstrated by a rightward shift of the response curves (Fig. 5). As the key interaction of our predicted binding mode, the mutation of D127A^3.32^, W286A^6.48^, and Y289A^6.51^ significantly reduced the G protein coupling ability, underscoring the essential role in odorant recognition of mTAAR7e. In the vicinity of Asp127^3.32^, Tyr316^7.42^ bridges with Asp127^3.32^ to stabilize the potential hydrogen-bond network, and a near complete abolishment of response was observed for mutant Y316A^7.42^ Additionally, varying degrees of reduction in activity were also observed in other amino acid mutations within the OBS, including V128A^3.33^, C131A^3.36^, S132A^3.37^, N217A^3.43^, and F290A^6.52^. Specifically, the polar interaction mutant N217A^5.43^ and the aromatic interaction mutant F290A^6.52^ show more pronounced effects, while the hydrophobic interaction mutant C131A^3.36^ exhibits a relatively weaker impact.

**Fig. 5.**
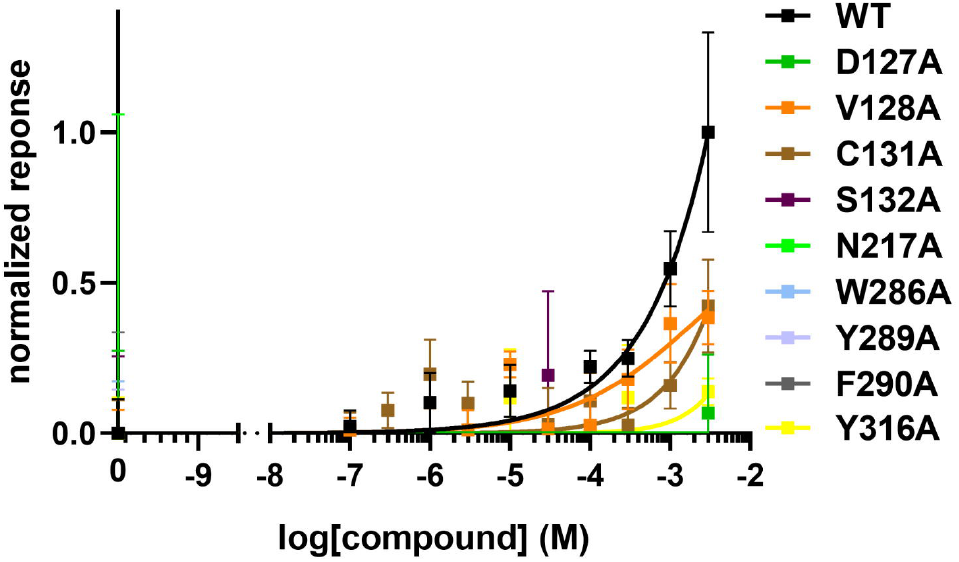
Responses of wild-type mTAAR7e and its binding site mutants on DMCHA. All responses are normalized to the highest response of the wild-type mTAAR7e to DMCHA. *N* = 4.

The response of the S132Y^3.37^ and S133C^3.38^ double mutant to DMCHA was greatly enhanced compared to the wild-type receptor (Fig. 6). Previous SAR studies on the experimental mTAAR7f indicated that odorant binding is highly dependent on the shape complementarity between the ligand and OBS, which is strongly affected by the CYC motif (Cys131^3.36^, Tyr132^3.37^, Cys133^3.38^) [5]. In the CYC motif, two cysteines form the disulfide bond to stabilize the shape of binding pocket, while the tyrosine residue separates the OBS into two distinct cavities. Unlike mTAAR7f, wild-type mTAAR7e harbors a CSS motif in place of CYC, eliminating the disulfide bond at positions 3.36/3.38 and the cavity partition. This yields a larger, more flexible pocket that enhances responsiveness to the long-chain ligand DMOA (8-carbon aliphatic chain). Conversely, mutating CSS to CYC restores sensitivity to DMCHA, which has a shorter fatty acid chain (Fig. 6). The same result was also found in DMBA. DMBA, which has a shorter chain length than DMOA, also showed increased binding activity to the doubly mutated mTAAR7e (SI Fig. S5). This finding suggests that the differences between the CYC and CSS motifs, through shaping the topological structure and stability of the binding pocket, are the main molecular driving force behind the distinct selective binding and activation of amine-type ligands of different sizes by mTAAR7e and mTAAR7f.

**Fig. 6.**
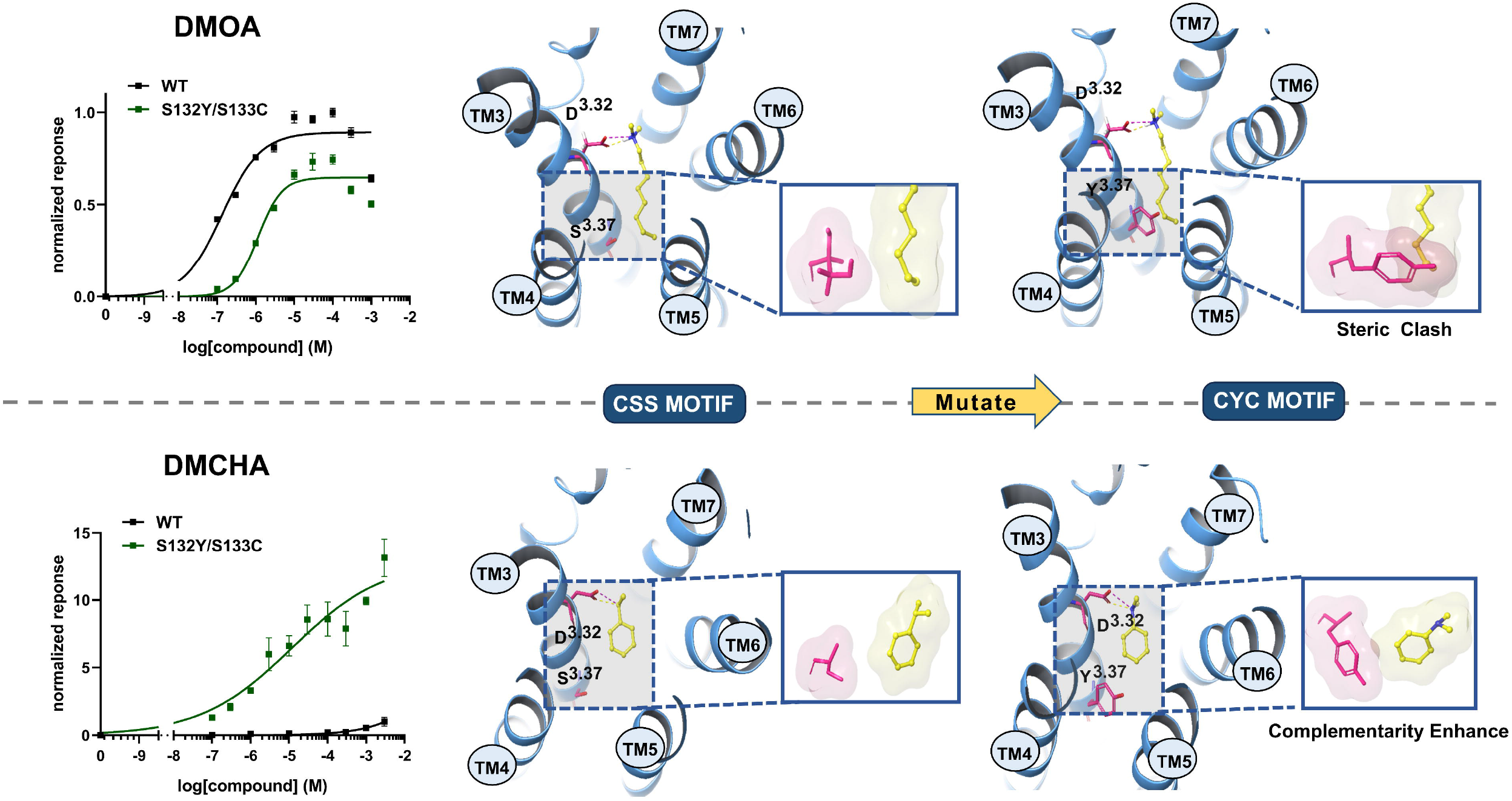
Responses and calculation analysis of wild-type mTAAR7e (CSS motif) and the S132Y/S133C double mutant (CYC motif) on DMOA and DMCHA. All responses are normalized to the highest response of the wild-type mTAAR7e to DMOA or DMCHA. *N* = 4.

### 3.4. Activation of mTAAR7e

The activation of class A GPCRs upon recognition of their respective ligands involves a series of structural changes, among which the most prominent is the outward shift of the intracellular C-terminal of TM6 enables coupling of G protein and the subsequent activation of signaling pathways [31, 32]. Two indicators are employed to quantify this outward displacement of TM6: TM6 distance and TM6 angle [33-35]. TM6 distance refers to the distance between the C_α_ atoms of residues 2.46 and 6.37, while TM6 angle represents the angle formed by the C_α_ atoms of residues 6.34, 6.47, and 2.41, with residue 6.47 serving as the apex of the angle. Statistical analysis of class A GPCR structures in different activation states reveals distinct distributions of TM6 distance and TM6 angle (Fig. 7b). During the inactive-to-active conformational transition, TM6 undergoes characteristic outward displacement, quantitatively reflected by increased values of both structural descriptors. To specifically analyze mTAAR7e activation, we aligned three distinct structural models: the apo state predicted by AlphaFold3 (AF3) [36], the HM structure, and a 300-ns MD simulation trajectory initiated from the HM structure (Fig. 7a). Compared to the AF3 model, the HM agonist-bound structure showed an outward shift of TM6, which is a hallmark of GPCR activation from inactive to active states. This conformational transition was validated by MD simulations, which confirmed the stability of these structural changes (Fig. 7b).

**Fig. 7.**
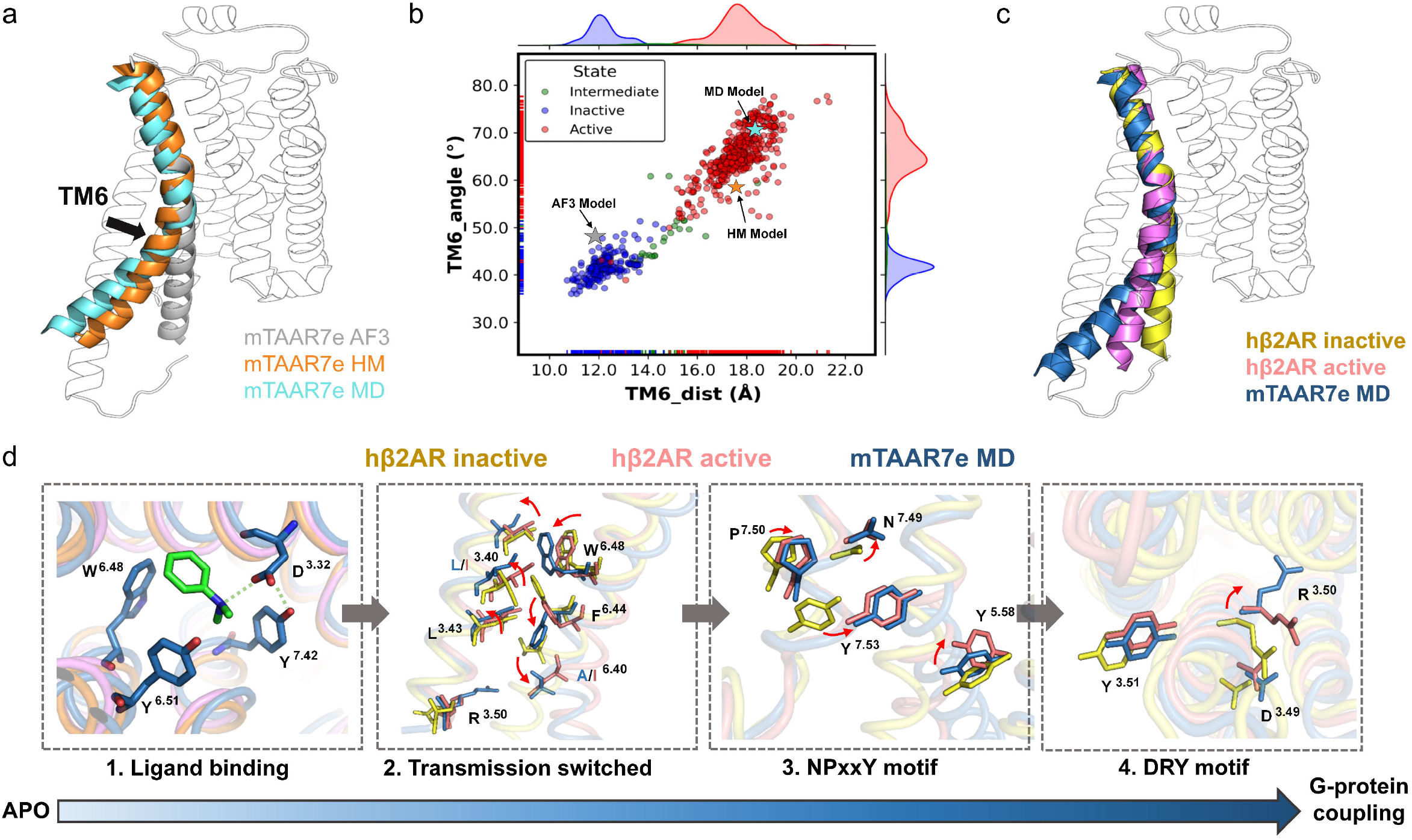
Conformational changes upon mTAAR7e activation. (a) Comparison of the 300-ns MD-simulated structure (cyan) of mTAAR7e, initially based on HM structure (orange), with the apo mTAAR7e model generated by AF3 (grey). (b) Scatter plot of TM6 distance versus TM6 angle in class A GPCRs. Data points are color-coded by GPCR activation state: active (red), intermediate (green), inactive (blue). Data markers are color-coded by mTAAR7e model source: AF3 (grey), MD simulation (cyan), HM (orange). (c) Global structural comparison of the inactive hβ_2_AR (red, PDB ID: 2RH1), active hβ_2_AR (violet, PDB ID: 3SN6), and mTAAR7e MD-simulated structure (slate). (d) Detailed alignment of functional motifs across inactive hβ_2_AR, active hβ_2_AR and mTAAR7e MD-simulated structure.

For further elaboration, we compared the structure of mTAAR7e with the experimentally resolved structures of the active and inactive states of hβ_2_AR (Fig. 7c) [37]. Residues critical for activation in mTAAR7e adopt positions analogous to those in the active hβ_2_AR, providing insights into its activation process. This process, broadly conserved among class A GPCRs, can be delineated into four sequential steps (Fig. 7d). The first step involves ligand recognition, the binding of DMCHA is primarily driven by electrostatic interactions between the tertiary amine of the ligand and Asp^3.32^, in addition to extensive van der Waals interactions in the hydrophobic pocket. Notably, the van der Waals interaction with Trp286^6.48^ induces an inward tilt (13°) compared to the inactive state of hβ_2_AR, and a similar but weaker tilt is also observed in the active state of hβ_2_AR. As the key element described in activation of many class A GPCRs, the shift of Trp286^6.48^ triggers propagation of structural changes in adjacent residues on TM6, including notable rotamer change of residue at positions Phe^6.44^ and Ala^6.40^, ultimately leads to the outward movement of the cytoplasmic end of TM6 [38]. As the increased distance between the C-terminal of TM3 and TM6, the spatial steric constraints on the conserved residue Arg^3.50^ at the C-terminal end of TM3 are alleviated, allowing it to adopt a low-energy conformation akin to that observed in the active state of hβ_2_AR. This conformational adjustment disrupts the ionic interaction between Asp^3.49^ and Arg^3.50^ within the activation motif DYR (Asp^3.49^ − Arg^3.50^ − Try^3.51^), a structural feature regarded as a hallmark of the inactive state in class A GPCRs [13]. Prior to the broken of the ionic lock in DYR motif, structural changes in the conserved motif NPxxY (Asn^7.49^ − Pro^7.50^ − x − x − Tyr^7.53^) on the C-terminal end of TM7 also contribute to the transition to the active state, where Asn^7.49^ contacts Pro^7.50^, facilitating a concerted stabilization across the sodium ion site and TM7 helix kink. The final amino acid in the NPxxY, a Tyr^7.53^ switch, make contacts with Tyr^5.58^, promoting the maintenance of the active state.

## 4. Conclusions

Since the initial discovery of TAARs, their ligand selectivity has been the subject of scrutiny. Past functional studies of TAARs uncovered structural characteristics required for receptor activation, including a clear separation in the ability to respond to amines of different degrees of substitution among the nine TAAR subfamilies. Our findings support that idea that key residues in the mTAARe binding pocket contribute to the preference for longer-chain, substituted amines. Given the olfactory and possible physiological roles of TAARs, the elucidation of the TAAR pharmacology will aid in the design and identification of new endogenous and exogenous ligands and the deorphaning of additional TAARs. While the activation mechanism for mTAAR7e proposed here remains a working hypothesis based on mutagenesis results and MD simulations, future studies aimed at determining the structures of additional mammalian TAARs may provide important insights on receptor function, regulation, and potential therapeutic targets.

## CRediT authorship contribution statement

**Yingjian Liu:** Writing – review & editing, Writing – original draft, Conceptualization, Supervision, Project administration. **Junhong He:** Writing – review & editing, Writing – original draft, Investigation, Formal analysis, Visualization. **Jiahui Sun:** Writing – review & editing, Writing – original draft, Investigation, Formal analysis, Visualization. **Chen Zhang:** Investigation, Formal analysis. **Peiei Shi:** Investigation, Formal analysis. **Hanyi Zhuang:** Writing – review & editing, Writing – original draft, Visualization. **Weihong Liu:** Writing – review & editing, Writing – original draft, Conceptualization, Investigation, Formal analysis, Supervision, Project administration.

## Declaraion of competing interest

The authors declare no conflict of interest.

## Appendix A. Supplementary data

Supplementary data to this article can be found online at http://XXX.

