## Supporting information for "Structural insights on the ligand selectivity of the mouse trace amine-associated receptor TAAR7e"

Supporting Figures and Tables

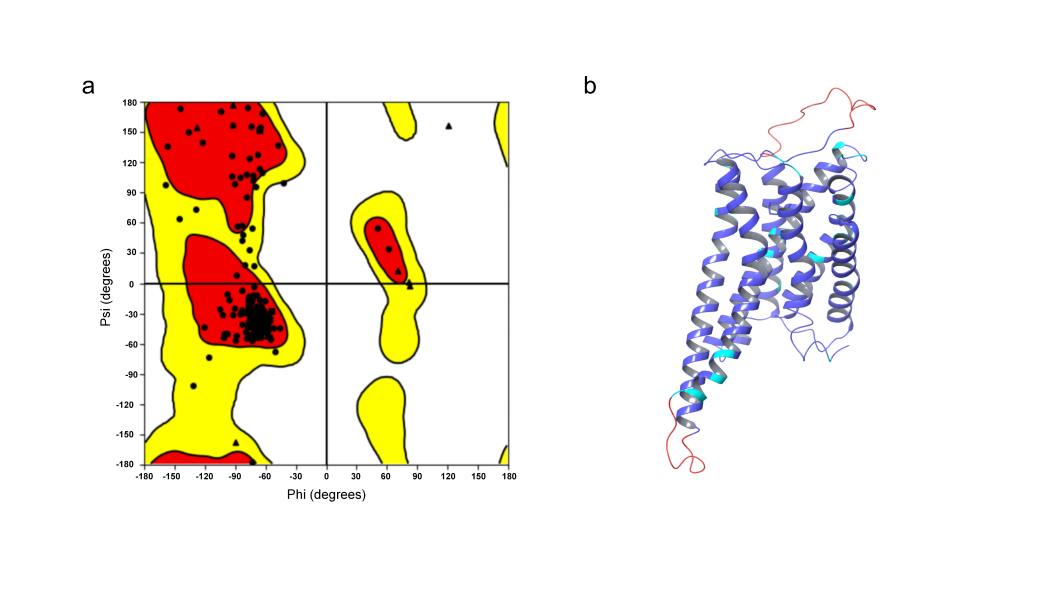

**Figure S1.** Assessment of the protein homology modeling structure for mTAAT7e. (a) Ramachandran plot of the mTAAT7e. (b) Confidence score of the mTAAT7e.

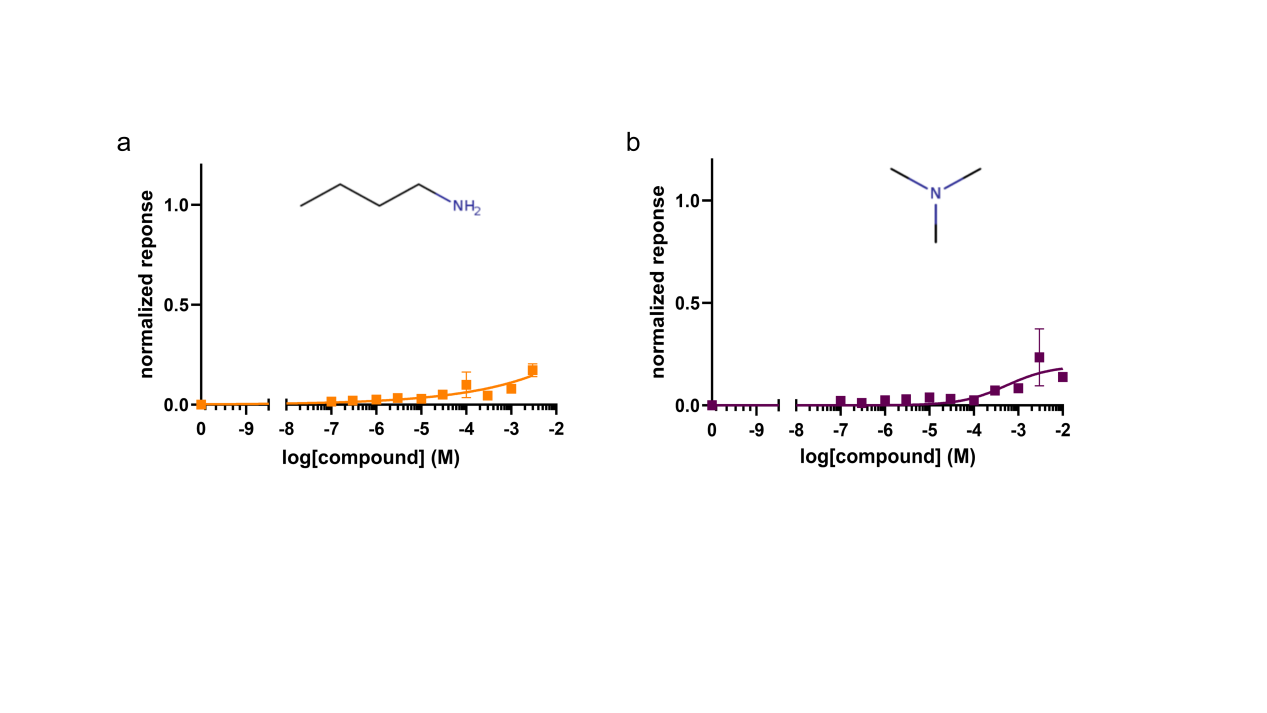

**Figure S2.** Response of mTAAR7e to Trimethylamine and Butylamine. All responses are normalized to the highest response to DMOA. *N* = 4.

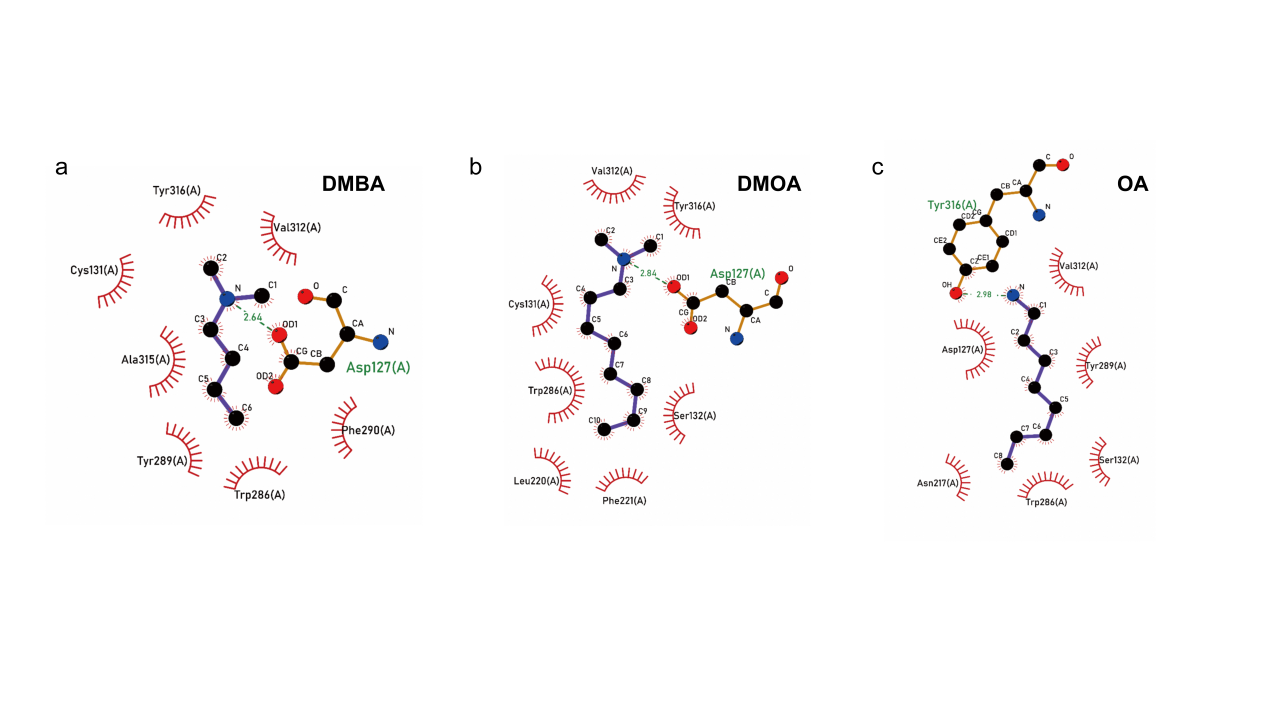

**Figure S3.** Ligand-mTAAR7e binding poses obtained from molecular docking. (a) DMBA. (b) DMOA. (c) OA.

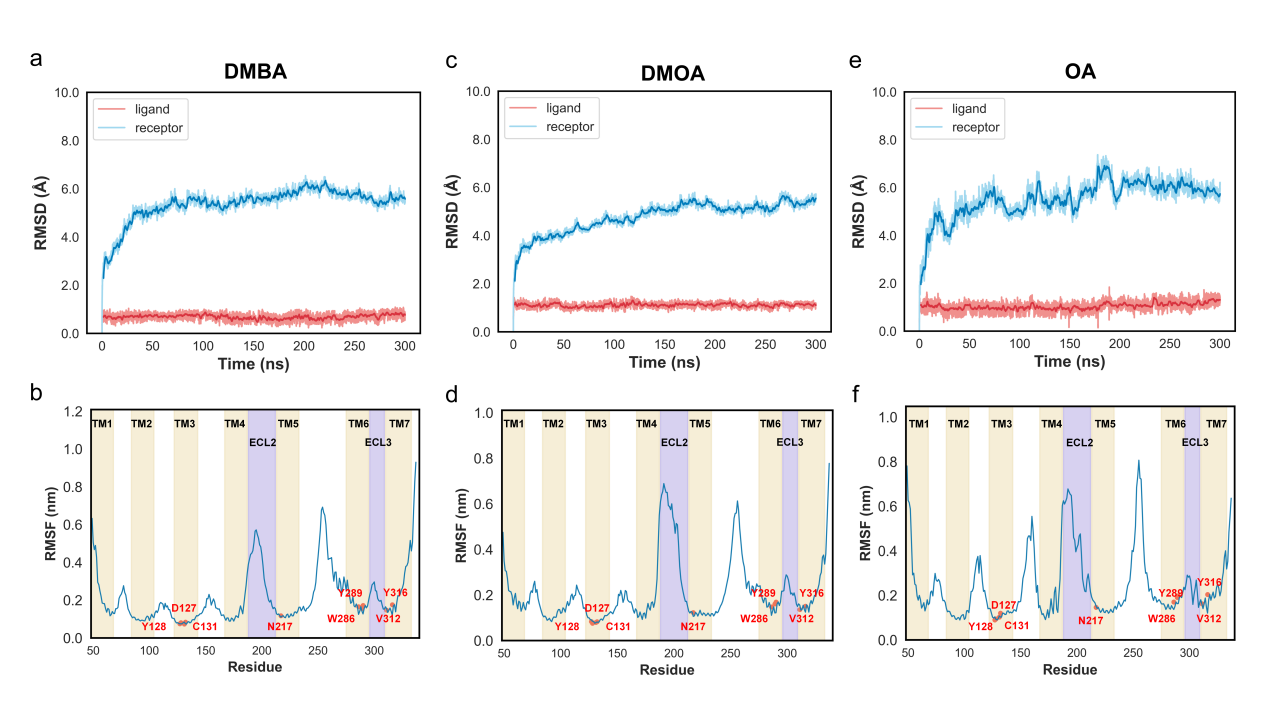

**Figure S4.** MD system stability. (a-b) RMSD and RMSF plots for DMBA. (c-d) RMSD and RMSF plots for DMOA. (e-f) RMSD and RMSF plots for OA.

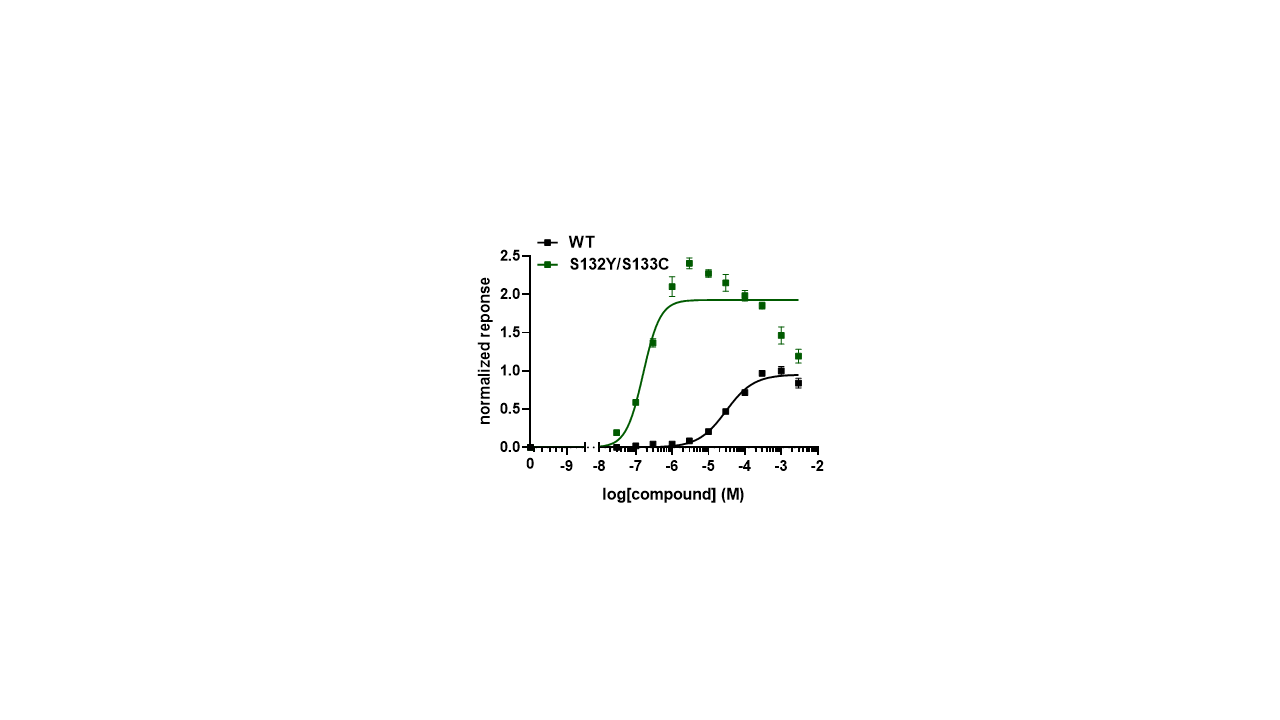

**Figure S5.** Response of wild-type mTAAR7e (CSS motif) and its mutant (CYC motif) on DMBA. All responses are normalized to the highest response of the wild-type mTAAR7e to DMBA. *N* = 4.

**Table S1.** Energy decomposition of four ligands from 3 rounds of 300-ns MD simulations via MMPBSA.

|  | **DMCHA** | **DMBA** | **DMOA** | **OA** |
| --- | --- | --- | --- | --- |
| **∆G_VDW_** | -17.96 ± 1.22 | -22.98 ± 0.74 | -34.96 ± 2.96 | -23.10 ± 0.62 |
| **∆G_EEL_** | -124.23 ± 3.64 | -129.42 ± 13.69 | -119.09 ± 10.54 | -94.94 ± 9.23 |
| **∆G_pb_** | 125.40 ± 1.52 | 122.40 ± 13.64 | 118.53 ± 9.15 | 98.92 ± 8.34 |
| **∆G_nonpolar_** | -16.08 ± 0.37 | -15.66 ± 0.51 | -24.00 ± 0.17 | -20.07 ± 0.46 |
| **∆G_disp_** | 25.11 ± 0.59 | 26.69 ± 1.27 | 40.11 ± 1.08 | 33.13 ± 0.68 |
| **∆G_gas_** | -142.19 ± 4.85 | -142.19 ± 13.19 | -154.05 ± 13.38 | -118.04 ± 8.62 |
| **∆G_solv_** | 134.44 ± 1.76 | 133.43 ± 12.63 | 134.64 ± 10.02 | 111.99 ± 8.17 |
| **∆G_total_** | -7.75 ± 3.34 | -18.96 ± 6.35 | -19.41 ± 3.41 | -6.06 ± 0.46 |
